# SWOTein: A structure-based approach to predict stability Strengths and Weaknesses of prOTEINs

**DOI:** 10.1101/2020.10.13.338046

**Authors:** Q. Hou, F. Pucci, F. Ancien, J.M. Kwasigroch, R. Bourgeas, M. Rooman

## Abstract

**Motivation:** Although structured proteins adopt their lowest free energy conformation in physiological conditions, the individual residues are generally not in their lowest free energy conformation. Residues that are stability weaknesses are often involved in functional regions, whereas stability strengths ensure local structural stability. The detection of strengths and weaknesses provides key information to guide protein engineering experiments aiming to modulate folding and various functional processes.

**Results:** We developed the SWOTein predictor which identifies strong and weak residues in proteins on the basis of three types of statistical energy functions describing local interactions along the chain, hydrophobic forces and tertiary interactions. The large-scale comparison of the different types of strengths and weaknesses showed their complementarity and the enhancement of the information they provide. We applied SWOTein to apocytochrome b_562_ and found good agreement between predicted strengths and weaknesses and native hydrogen exchange data. Its application to an amino acid-binding protein identified the hinge at the basis of the conformational change. SWOTein is both fast and accurate and can be applied at small and large scale to analyze and modulate folding and molecular recognition processes.

**Availability:** The SWOTein webserver provides the list of predicted strengths and weaknesses and a protein structure visualization tool that facilitates the interpretation of the predictions. It is freely available for academic use at http://babylone.ulb.ac.be/SWOTein.

## 1 Introduction

The class of structured proteins have to adopt a well-defined 3-dimensional (3D) structure in order to be functional. Even though this folded structure generally corresponds to the global free energy minimum in physiological conditions, most interactions taken individually are not in their minimum free energy state. Indeed, the chemical moieties that make up the proteins are not free to move but are constrained within the amino acids, which are in turn constrained within the polypeptide chain. These constraints are often incompatible with lowest free energy amino acid interactions and conformations. They add a level of difficulty in unraveling the protein folding process.

The kinetics of protein folding is another crucial factor. Natural proteins must fold fast enough – in times of the order of the second – to be able to reach their functional state in the crowded cell environment or inside chaperone protein chambers. This imposes the free energy landscape to be smooth enough to prevent proteins from getting stuck in misfolded states.

The optimal satisfaction of this ensemble of thermodynamic and kinetic constraints is known as the minimal frustration principle [1, 2, 3]. Natural protein sequences have evolved to satisfy this principle to a certain extent [4], but functional constraints prevent complete optimization. Indeed, a protein must not only fold fast and be stable, it has also to be functional. Proteins usually contain low-frustration regions that are well optimized for stability, which we call stability strengths, and frustrated regions that are not optimized for stability, which we refer to as stability weaknesses. Note that stability is defined in terms of free energy, and thus incorporates both enthalpic and entropic contributions.

Several measures of frustration have been designed. Mutational frustration compares the energy of each wildtype contact with the energy of the same contact formed by mutant residues. It thus integrates the effect of natural evolution [5, 6, 7, 8]. Similarly, stability strengths and weaknesses are identified as residues that, when mutated, lead on the average to strong destabilization or strong stabilization, respectively [9, 10], as estimated with the PoPMuSiC algorithm [11, 12, 10].

Configurational frustration is a closely related concept defined in structure space rather than in sequence space. It compares the strength of the interactions in the native structure and in decoy structures [5, 6, 7, 8], in alternative non-native structures [9, 13, 14, 15], or in structures modeling the unfolded state [16].

It is the latter definition of stability strengths and weaknesses that we considered here. We estimated in a recent study [16] the residue contributions to the global folding free energy through three types of database-derived statistical potentials based on three kinds of structure elements, *i.e*. inter-residue distances, backbone torsion angles and solvent accessibility. Very positive folding free energy contributions were taken to indicate stability weaknesses and very negative folding free energy contributions, stability strengths. This approach was successfully applied to reveal the structural and functional characteristics of bovine seminal ribonuclease [16].

Here, we further improved our approach to identify stability strengths and weaknesses, performed a series of large-scale analyses, compared our predictions with experimental hydrogen-exchange data, applied our algorithm to three case studies, and provided a webserver for non-expert users, called SWOTein.

## 2 Methods

### 2.1 A brief summary of statistical potentials

The formalism of statistical potentials [17, 18, 19, 20] has long been used in protein science to study biophysical protein properties, which range from mutational robustness to thermal resistance and protein-protein interactions. These potentials are knowledge-based mean-force potentials derived from datasets of experimentally resolved 3D protein structures using the inverse Boltzmann law. In a nutshell, this relation associates the free energy of a state to the probability of observing the system in that state.

The statistical potentials that we used here are based on a coarse grained representation of protein structures, in which only the heavy main chain atoms and the amino acid-dependent average geometric center of the side chains were considered. Explicit side chain degrees of freedom were thus overlooked.

Let us suppose that *C* is a conformational descriptor such as the distance between two residues, and *S* a sequence descriptor such as an amino acid type or a pair of amino acids. The folding free energy associated to the sequence-structure association (*S, C*) is defined in terms of the conditional probabilities *P*(*C*|*S*) and *P*(*C*|*S*)_ref_ of observing *C* given *S* in the native state and in a reference state, respectively:

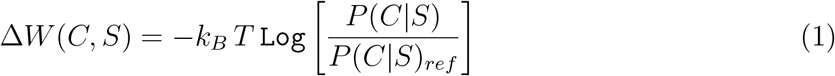

where *k_B_* is the Boltzmann constant and *T* the absolute temperature. Here we chose the reference state as the state for which the probability to observe a given conformation is independent of the sequence:

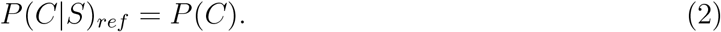

This reference state mimics the unfolded state or an average misfolded state.

The folding free energy is thus finally expressed as:

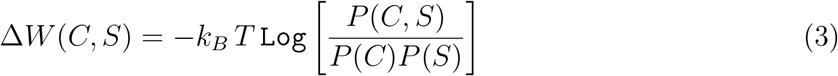

The probabilities appearing in this equation are estimated from the frequencies of observation of the sequence and structure elements *S* and *C* in a good-quality and non-redundant dataset of experimental 3D protein structures, in the approximation that this set is large enough to yield statistically significant sequence and structure sampling. In terms of the number of occurrences *n_S_*, *n_C_* and *n_cs_* of *S* and/or *C* in the dataset and their total number *n*, the folding free energy reads as:

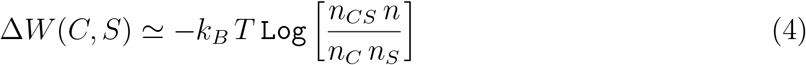

SWOTein uses three different statistical potentials, constructed from three different conformational descriptors *C*, described below.

#### Solvent accessibility potential Δ*W_acc_*

This potential describes the interaction of protein residues with the solvent [21, 22]. The conformational descriptor *C* is here the solvent accessibility (*a*_1_,*a*_2_) of two different residues, and the sequence descriptor *S* is an amino acid type *s*. (*a*_1_, *a*_2_) and *S* are situated in a sequence window of five residues. The solvent accessibility of a residue is defined as the ratio of its solvent accessible surface area in a given structure and in an extended Gly-X-Gly tripeptide conformation, and is computed with our in-house software implementation [23]. It ranges from 0% when the residue is completely buried to 100% when it is fully exposed to the solvent. This interval is split into five bins (0-5%, 5-15%, 15-30%, 30-50%, 50-100%). For this potential, Eq. (4) reads as:

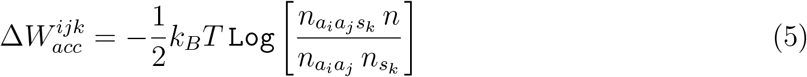

where *i, j* and *k* represent residue positions in a sequence window of five consecutive residues, with *i* ≠ *j*. More precisely, *s_k_* is the amino acid type at position *k* along the sequence, and *a_i_* and *a_j_* are the solvent accessibility bins of the *i*th and *j*th residues along the sequence.

#### Torsion angle potential Δ*W_tor_*

This potential describes the local interactions along the polypeptide chain in terms of the propensities of amino acids to have their (*φ, ψ, ω*) backbone torsion angles falling in one of the seven domains defined in [24]. The conformational descriptor *C* is here the *φ, ψ, ω* domain of two different residues, noted (*t*_i_, *t*_2_), and the sequence descriptor *S* is an amino acid type *s*. For this potential, Eq. (4) reads as:

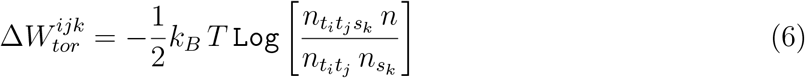

where *i, j* and *k* represent residue positions along the chain in a sequence window of seven consecutive residues, with *i ≠ j*.

#### Distance potential Δ*W_dis_*

The last potential describes tertiary interactions and is based on spatial distances *d* between protein residues [21, 22]. These distances were computed between the average geometrical centers of heavy side chain atoms and range between 3.0 to 8.0 Å. This interval is split into 25 bins of 0.2 Å width, with two additional bins grouping the distances that are smaller than 3 Å and larger than 8.0 Å, respectively. For the distance potential, Eq. (4) reads as:

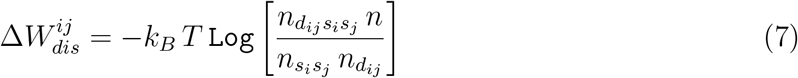

where *i* and *j* are sequence positions separated by one residue at least.

### 2.2 Protein structure dataset 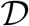

The dataset of protein structures used in the derivation of the statistical potentials consist of X-ray structures collected from the Protein Data Bank (PDB) [25], with a resolution of at most 2.0 Å. We considered only monomeric structures as annotated by the crystallographers who solved the structure or, if unavailable, by the PISA webserver [26]. We then used the culling webserver PISCES [27] to select the structures that share a maximum pairwise sequence identity of 20%. In this way, we obtained our dataset 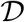 of 3,005 high-quality and non-redundant protein structures.

The three statistical potentials defined in Eqs (5)–(7) and derived from the dataset 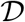 are explicitly given in the GitHub repository https://github.com/3BioCompBio/SWOTein, with technical details and differences with SWOTein’s preliminary version [16]. Note that the sizes of the sequence windows appearing in the definition of the potentials, Eqs (5)–(7), have been selected to obtain the lowest average folding free energy values on the dataset 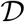.

## 3 Results

### 3.1 Strengths and weaknesses

To define strengths and weaknesses in a protein structure, we estimated the contribution of each residue in a given protein to the overall folding free energy. For that purpose, we considered all contributions Δ*W*(*C, S*) defined in Eq. (3) and divided them equally among the structural states that are contained in *C* [16]. More specifically, for the accessibility, torsion and distance potentials defined in Eqs (5), (6), (7), we allocated an equal folding free energy contribution to each of the two residues *i* and *j* that carry the structural state *C*, as exemplified in Fig. 1. This yields, for each of the three potentials:

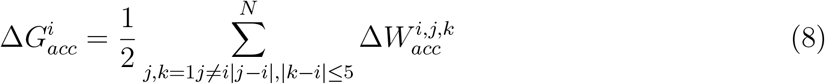

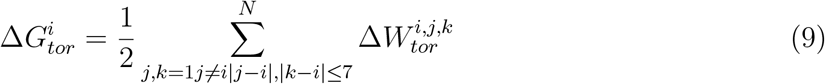

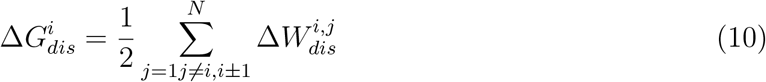

where *N* is the sequence length. Since no *φ*-angle is defined for the first residue of a protein chain and no *ψ*-angle for the last, no torsion folding free energy contribution 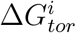 can be associated to these residues. An identical problem arises if there is a chain break or if some residues or relevant atoms are missing.

**Figure 1:**
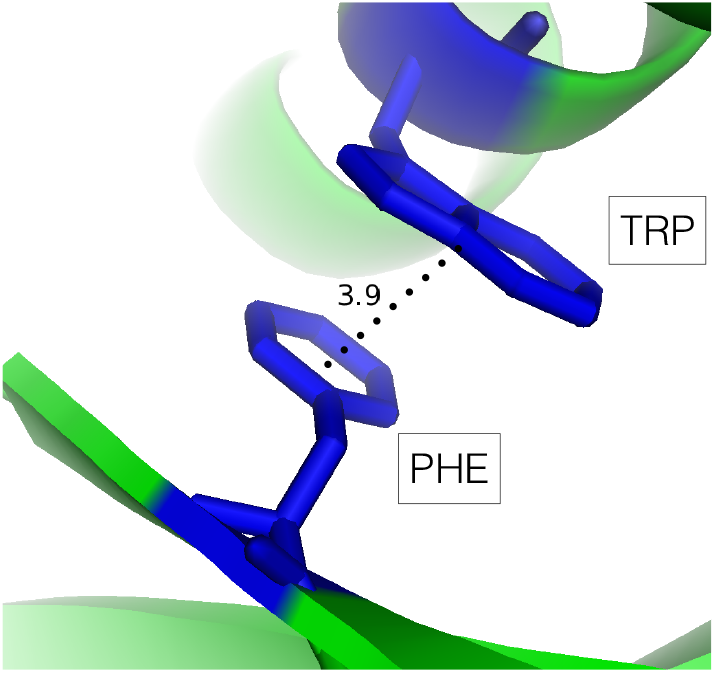
Division of the interaction energy between a Phe-Trp interacting pair at a spatial distance of 3.9 Å: one half of 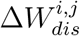 is attributed to the Phe residue at position *i* (on 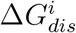) and one half to the Trp residue at position *j* (on 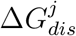).

We computed the distributions of folding free energy values 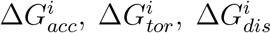 for all residues of all proteins in the dataset 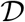. They are shown in Fig. 2, and their mean and standard deviation are given in Table 1. The values of the mean are all negative, indicating that most residues have stabilizing contributions. The left standard deviation *σ*^-^ is higher than the right standard deviation σ^+^ for 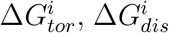, whereas the opposite is true for 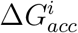.

**Figure 2:**
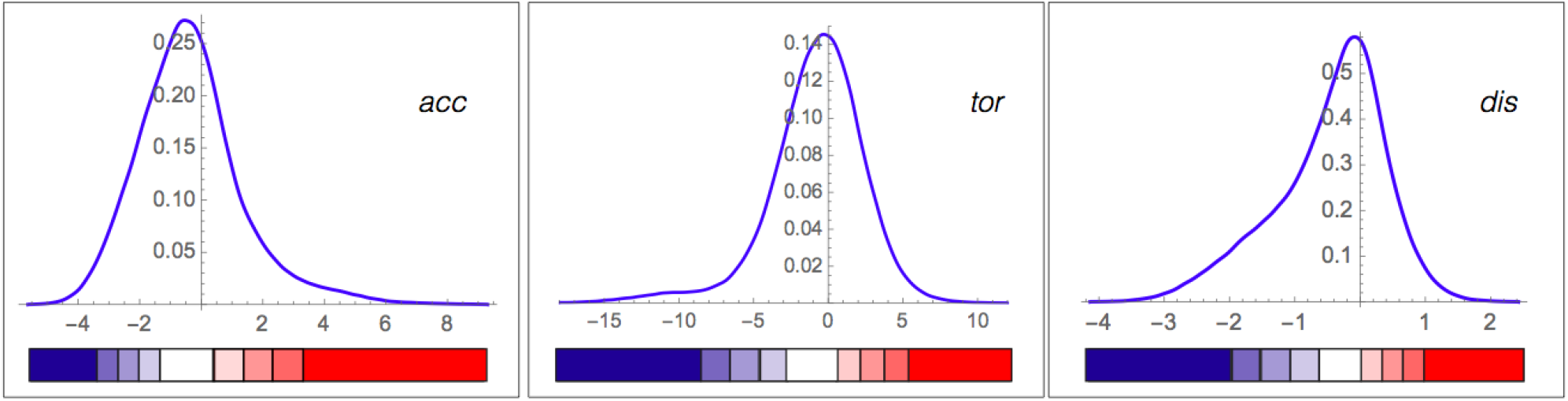
Distributions of per-residue folding free energy contributions (in kcal/mol) 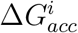, 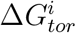 and 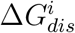 for all residues belonging to 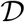. The classes of strengths and weaknesses are shown below the graphs, using the color code defined in Table 2.

**Table 1:**
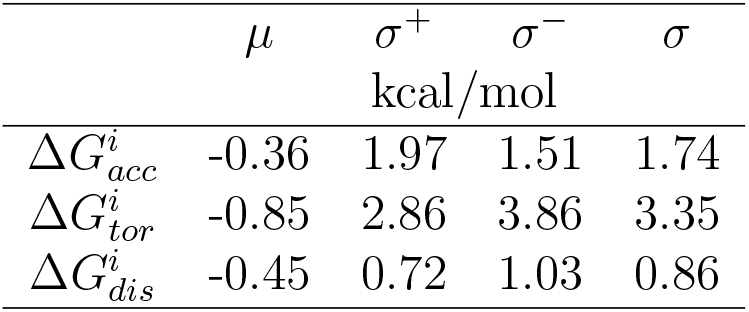
Folding free energy distribution for all residues belonging to 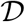: mean *μ*, standard deviation σ, left standard deviation σ^-^ computed from the folding free energy contributions that are lower than the mean, and standard deviation σ^+^ computed from the folding free energy contributions that are larger than the mean.

| | $\mu$ | $\sigma^+$ | $\sigma^-$ | $\sigma$ |
| --- | --- | --- | --- | --- |
|  | kcal/mol |  |  |  |
| $\Delta G_{acc}^i$ | -0.36 | 1.97 | 1.51 | 1.74 |
| $\Delta G_{tor}^i$ | -0.85 | 2.86 | 3.86 | 3.35 |
| $\Delta G_{dis}^i$ | -0.45 | 0.72 | 1.03 | 0.86 |

We defined different degrees of stability strengths and weaknesses on the basis of the values of 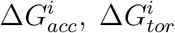 and 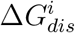 with respect to threshold values that depend on the mean and standard deviation of the folding free energy distributions, as reported in Table 2. More specifically, we have defined nine subclasses: hard strength, strength, moderate strength, mild strength, neutral, mild weakness, moderate weakness, weakness and hard weakness.

**Table 2:**
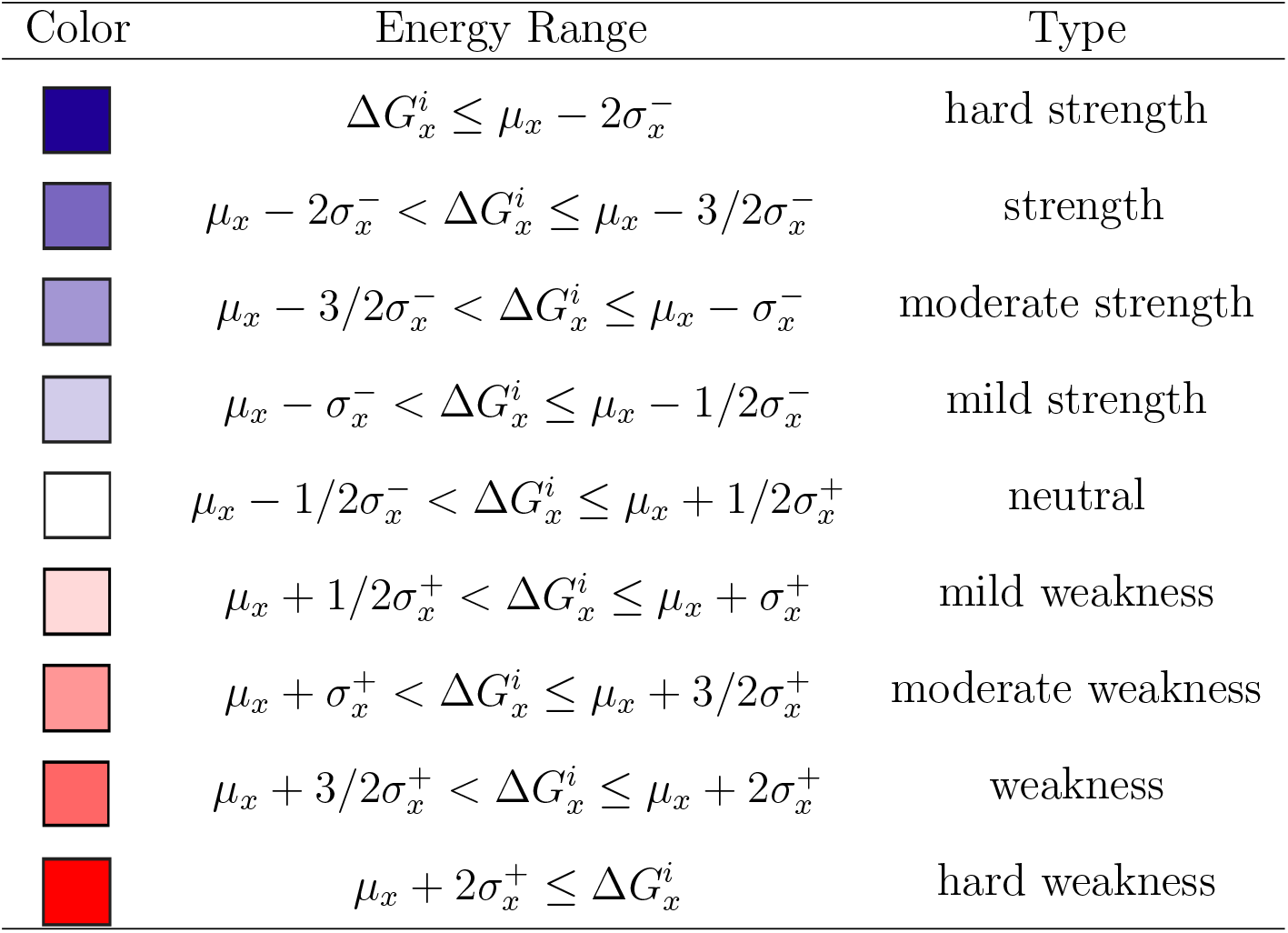
Definition of classes of stability strengths and weaknesses in terms of the mean and standard deviations given in Table 1, for each of the three folding free energy contributions 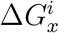 where ‘x‘ corresponds to ’dis’, ’acc’ or ’tor’. The colors used in Fig. 2 and in the visualization framework of the SWOTein webserver are here defined.

| Color | Energy Range | Type |
| --- | --- | --- |
| | $\Delta G_x^i \leq \mu_x - 2\sigma_x^-$ | hard strength |
| | $\mu_x - 2\sigma_x^- < \Delta G_x^i \leq \mu_x - 3/2\sigma_x^-$ | strength |
| | $\mu_x - 3/2\sigma_x^- < \Delta G_x^i \leq \mu_x - \sigma_x^-$ | moderate strength |
| | $\mu_x - \sigma_x^- < \Delta G_x^i \leq \mu_x - 1/2\sigma_x^-$ | mild strength |
| | $\mu_x - 1/2\sigma_x^- < \Delta G_x^i \leq \mu_x + 1/2\sigma_x^+$ | neutral |
| | $\mu_x + 1/2\sigma_x^+ < \Delta G_x^i \leq \mu_x + \sigma_x^+$ | mild weakness |
| | $\mu_x + \sigma_x^+ < \Delta G_x^i \leq \mu_x + 3/2\sigma_x^+$ | moderate weakness |
| | $\mu_x + 3/2\sigma_x^+ < \Delta G_x^i \leq \mu_x + 2\sigma_x^+$ | weakness |
| | $\mu_x + 2\sigma_x^+ \leq \Delta G_x^i$ | hard weakness |

### 3.2 Large-scale analysis

It is interesting to see how strengths and weaknesses depend on the solvent accessibility 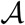. We plotted the stability contributions 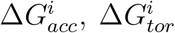 and 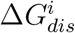 as a function of the solvent accessibility of the residues *i*. As shown in Fig. 3, we observed different behaviors according to the potentials used for identifying the strengths and weaknesses. Using the solvent accessibility potential, the strong and weak residues are preferentially located in the core with 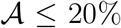, or in a partially buried area with 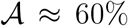. In contrast, the number of strong and weak residues is basically constant when using the torsion potential, whereas it decreases almost linearly from the core to the surface when using the distance potential. This result is related to the fact that tertiary inter-residue interactions described by the distance potential contribute more to the overall protein stability when situated in the core, whereas the local interactions along the chain described by torsion potentials are equally important in the core and on the surface [28]. The accessibility potential has a mixed behavior, between that of tertiary and local interactions.

**Figure 3:**
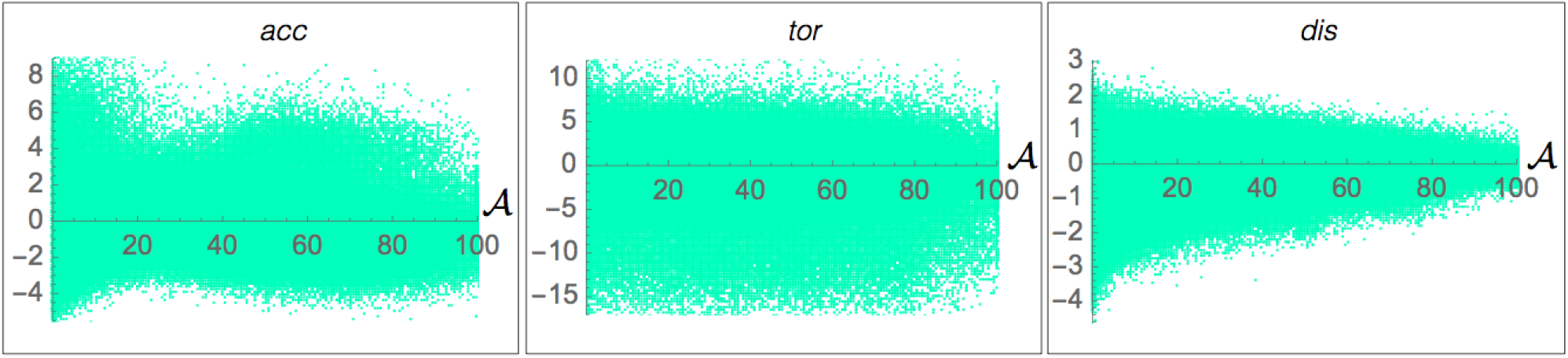
Folding free energy contributions 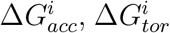 and 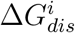 (in kcal/mol) as a function of the solvent accessibility 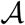, for all residues *i* belonging to 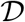.

We examined whether a strength identified by one potential is also a strength for the other potentials, and similarly for the weaknesses. As shown in Fig. 4, it is absolutely not the case. Especially the torsion potential gives very different information: it is neither correlated with the solvent accessibility potential nor with the distance potential; the Pearson correlation coefficients are indeed as low as 0.07 and 0.04, respectively. The solvent accessibility and distance potentials are slightly better correlated, with a correlation coefficient of 0.38. These results are not surprising as the torsion potential describes purely local interactions along the chain, whereas the distance potential and the solvent accessibility potential describe tertiary residue-residue and residue-solvent interactions, respectively, which are dominated by the hydrophobic effect [28, 21].

**Figure 4:**
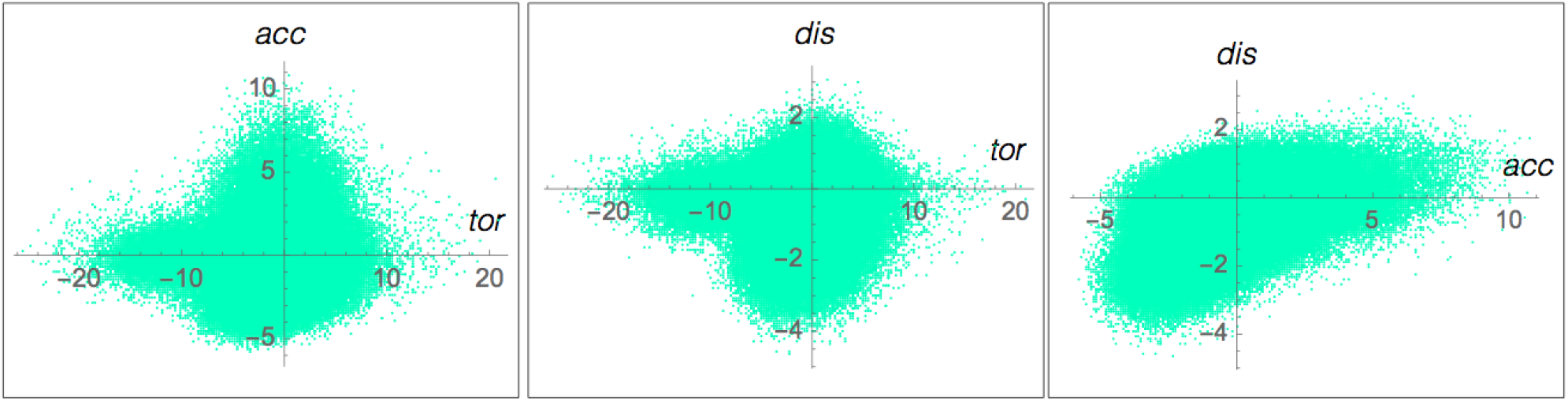
Pairwise comparison between the per-residue folding free energy contributions 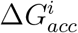, 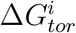 and 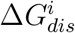 (in kcal/mol), for all residues *i* belonging to 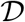.

The use of several potentials describing different types of interactions is a unique characteristic of our approach: it yields multiple, complementary, indicators of stability strengths and weaknesses, and is thus more informative than methods based on a single value.

### 3.3 Comparison with experimental stability values

The level of stability of protein regions can be experimentally determined by using hydrogen exchange techniques [29]. These techniques are based on the property of amide hydrogens that are (transiently or permanently) exposed to the solvent to exchange with the hydrogens of the solvent. The more the exchange between amide protons and solvent protons is retarded upon unfolding, the more stable the structure in which the amide protons are involved [30]. By measuring how the free energy of hydrogen exchange (Δ*G_HX_*) of each residue varies with denaturant concentration, it is possible to determine which protein regions totally unfold, partially unfold or remain native-like at low denaturant concentration. These regions correspond to what we call weak, intermediate and strong regions, respectively.

We collected curated hydrogen exchange data from the Start2Fold database [31], which contains 57 proteins with experimental 3D structure, wherein 281 residues are classified as weak, 628 as medium and 1,088 as strong.

For each of these residues, we computed the per-residue folding free energy contributions 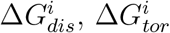 and 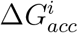. As shown in Fig. 5, the folding free energy is on the average much lower for the residues experimentally shown to be strong, higher for the medium residues and even much higher for the weak residues. This is true for the three potentials, which leads us to conclude that they correctly capture the local stability features of protein structures. Note that experimentally identified weak residues are not always weak for the three potentials, and similarly for the strong residues. As we will see in the case studies presented in the next sections, this is particularly interesting as it gives information on the type of stabilizing forces involved.

**Figure 5:**
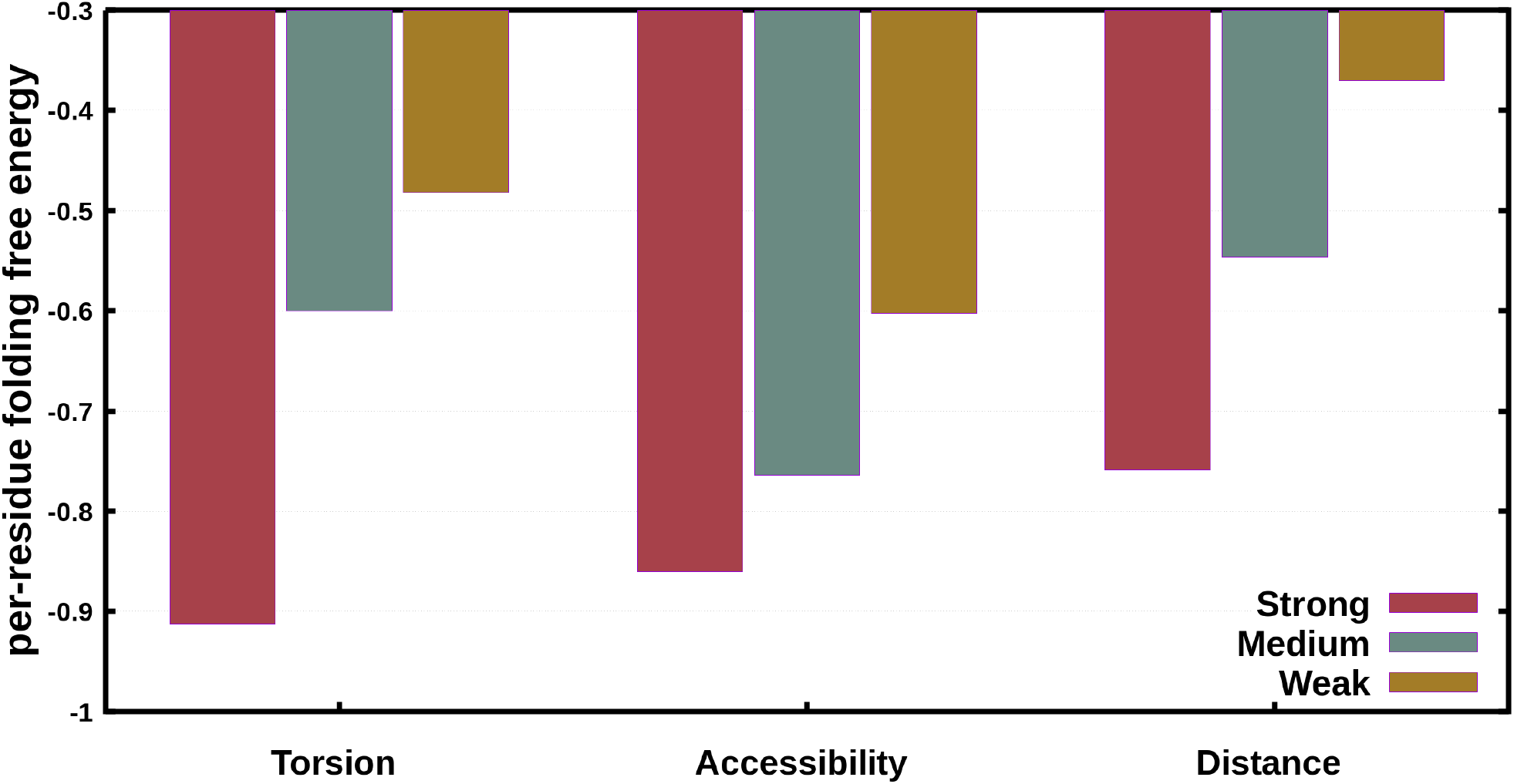
Comparison between experimental and computed strengths and weaknesses. Average per-residue folding free 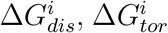 and 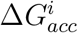 (in kcal/mol) for the residues *i* experimentally identified as strong, medium and weak in [31]. The average folding free energy differences between weak/medium, weak/strong and medium/strong sites are, for the majority, statistically significant for the three potentials; details are given in Supplementary Material.

The comparison between the prediction quality of SWOTein, its preliminary version [16] and the Frustratometer [7] in distinguishing between weak, medium and strong sites is shown in Supplementary Material, with an incontestable advantage for SWOTein.

### 3.4 SWOTein webserver

The SWOTein program runs on a user-friendly and freely available webserver at http://babylone.ulb.ac.be/SWOTein/index.php. It automatically identifies strengths and weaknesses in experimental or modeled 3D structures of target proteins. The user can either provide the 4-letter code of a structure available in the Protein Data Bank (PDB) [25], which is then automatically retrieved, or upload a personal structure file in PDB format. After this step, the user chooses the chain(s) to analyze, submits the query and the computational process starts.

Note that selecting a subset of the chains in the PDB file amounts to completely discard the non-selected chains. This has an impact on the accessibility potential as the solvent accessibility is computed by considering only the selected chains. It has also an impact on the distance potential as interactions between residues located in the selected and non-selected chains are not considered.

SWOtein then provides a job ID and a link that must be followed or bookmarked to check the job’s status. When the computation is finished, the results are given on the same page. The computation is usually very fast: the analysis of a medium-size protein takes about 30 sec.

The webserver outputs different results upon completion of the computations, among which a table containing the values of 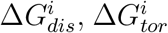 and 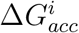 for each residue *i* in the target protein. Each folding free energy contribution is associated to a class and to a color that ranges from blue (hard strength) to red (hard weakness), with all intermediate color graduations defined in Table 2. These colors are also shown on the three-dimensional structure of the target protein (following the link “visualization tool”). Three different views are provided, one for each potential, given that the strengths and weaknesses differ according to the potentials.

The webserver also provides a downloadable text file containing the folding free energy contributions computed by the three potentials, associated to all residues in the target protein. More information and explanations can be found on the webserver’s help page.

### 3.5 Application of SWOTein

We present three case studies showing how the SWOTein webserver can be used to identify strengths and weaknesses in proteins of interest. We already used a first version of our method to analyze bovine seminal ribonuclease [16]. Here we applied SWOTein to investigate the structural properties of a redesigned apocytochrome b_562_, an amino acid-binding protein, and a subtilisin-eglin c complex.

It has to be emphasized that SWOTein predictions can be used to identify relevant residues to mutate in order to modulate the folding process and to stabilize or destabilize the native structure or protein-protein complex. SWOTein is thus very helpful to rationally guide protein engineering experiments.

#### Redesigned apocytochrome b_562_ (Rd-apocyt b562)

We considered Rd-apocyt b_562_, a redesigned four-helix bundle protein in which the residues around the heme present in the holo form were substituted with hydrophobic residues. The structure of this protein has been solved by nucleic magnetic resonance spectroscopy (NMR) and contains 15 models; its PDB code is 1YYX.

In order to experimentally study the presence of partially unfolded forms of Rd-apocyt b_562_, native-state hydrogen-exchange experiments have been performed [32], which identified four structural units with decreasing degrees of stability. Helix II, the C-terminal part of helix III and the N-terminal region of helix IV (residues 26-40 and 70-93) belong to unit 1, the C-terminal end of helix IV (residues 70-93) to unit 2, helix I (residues 1-25) to unit 3, and the loop between helices II and III and the N-terminal part of helix III (residues 41-69) to unit 4 [32].

The SWOTein predictions agree well with the stability of these units, as seen in Table 3 and Fig. 6. We computed the per-residue folding free energy contributions using the three potentials, 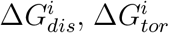 and 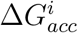, for the 15 NMR models contained in the PDB file 1YYX. We averaged these contributions over the 15 models, and over the residues contained in the four structural units of decreasing stability (Table 3). We found a very good agreement between the measured stability of the units and the average free folding energy contributions computed with the torsion angle potential, the distance potential and the sum of the three potentials, *i.e*. 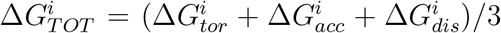. Indeed, the Spearman’s rank correlation coefficients are equal to 0.97 (p-value 10.005), 0.86 (p-value ~ 0.06) and 0.97 (p-value 10.005), respectively. The torsion angle potential is particularly interesting: the 2 subparts of unit 1 are strengths, units 2 and 3 are neutral, and unit 4 is a weakness (Table 3). This indicates the important role of local interactions along the chain in the (un)folding process of this protein. In contrast, the solvent accessibility contributions are poorly correlated, thus suggesting the lower role of hydrophobic forces in the process. The different performance of the three potentials is not a general result but is specific to Rd-apocyt b_562_. It indicates that the considered protein folds more according to a hierarchical folding model than to a hydrophobic collapse model, in other words, that the local, secondary, structure formation basically precedes the collapse into a compact globule [33, 34].

**Table 3.** Computed folding free energy contributions (in kcal/mol) for Rd-apocytb 562 (1YYX), averaged over the residues involved in the structural units identified in [32] and over the 15 NMR models in 1YYX. Unit 1 is separated into two subunits containing consecutive residues, which have the same measured stability. The color code is defined in Table 2.

| Structural unit | Residue range | $\langle \Delta G_{acc}^i \rangle$ | $\langle \Delta G_{tor}^i \rangle$ | $\langle \Delta G_{dis}^i \rangle$ | $\langle \Delta G_{TOT}^i \rangle$ |
| --- | --- | --- | --- | --- | --- |
| kcal/mol |  |  |  |  |  |
| unit 1 | 26-40 | -2.0 | -5.9 | -0.5 | -2.8 |
| unit 1 | 70-93 | -1.3 | -4.2 | -0.4 | -2.0 |
| unit 2 | 94-105 | 0.7 | -1.7 | 0.0 | -0.3 |
| unit 3 | 1-25 | -0.1 | 0.3 | 0.0 | 0.1 |
| unit 4 | 41-69 | -0.5 | 3.2 | 0.0 | 0.9 |

**Figure 6.**
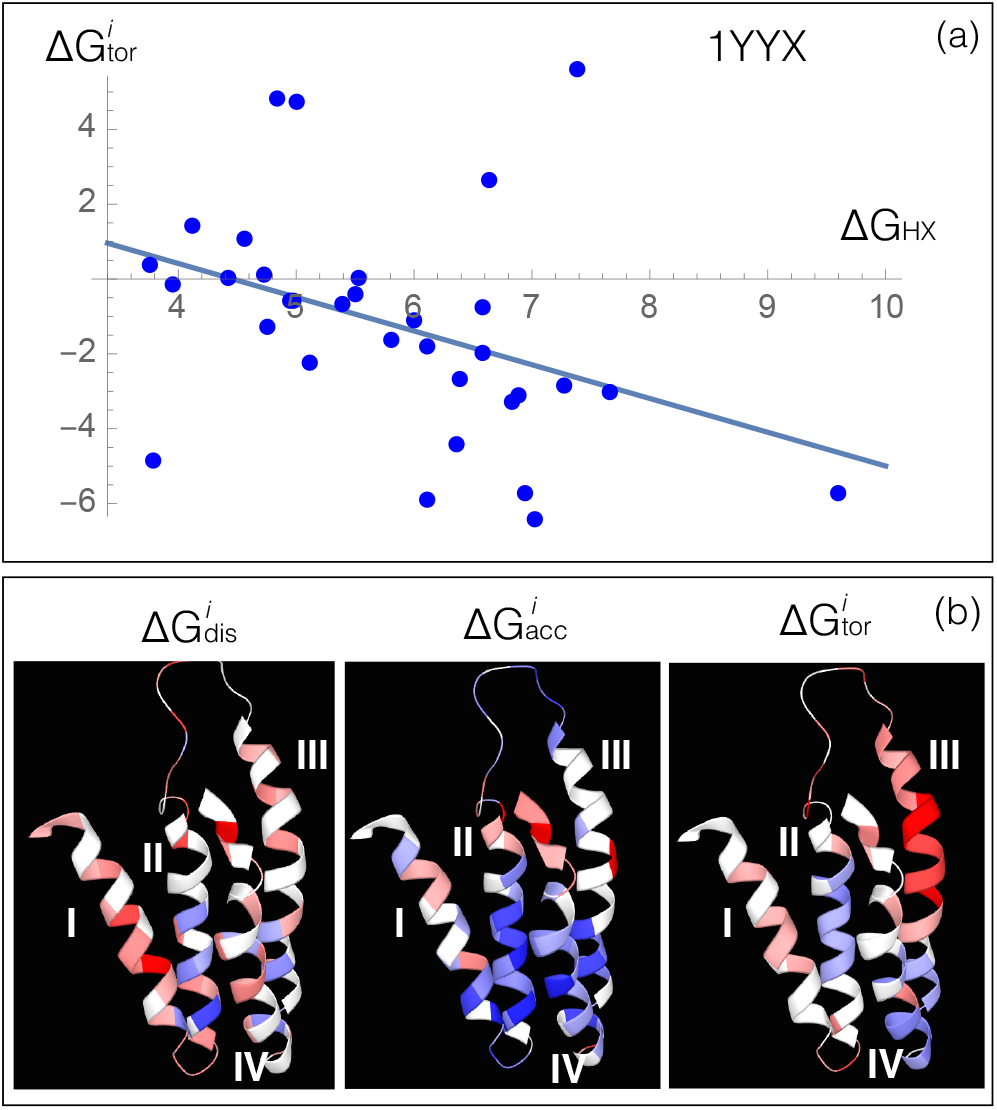
Per-residue folding free energy contributions in Rd-apocytb 562 (1YYX) (a) Folding free energy contributions 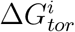 (in kcal/mol) averaged over the 15 NMR models contained in 1YYX as a function the experimental hydrogen exchange free energy Δ*G_NH_*, for all the residues for which the latter has been measured in [32]. (b) Visualization of the predicted strengths and weaknesses obtained from the three potentials, using the first NMR model in 1YYX and the color code defined in Table 2. These figures are directly taken from the SWOTein webserver.

Let us now focus on the per-residue folding free energy contributions rather than on the average values of the folding units. The good predictive power of the torsion potential in the stability Rd-apocyt b_562_ is also visible in Fig. 6.a, where the per-residue contributions 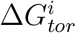, averaged over the 15 NMR models, are plotted as a function of the experimental free energy values of hydrogen exchange Δ*G_HX_*. The Pearson correlation coefficient is equal to −0.39, which is a very good result considering that we did not perform any parameter fitting. The 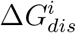 correlation is also high (r=-0.37), while 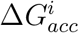 correlates only poorly (r=-0.05). The total free energy value 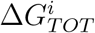 has a correlation coefficient of −0.39.

The per-residue folding free energy contributions for the first NMR model in the PDB structure 1YYX are shown in Fig. 6.b, for the three potentials. These figures are directly outputted from the SWOTein webserver.

#### Lysine/arginine/ornithine-binding protein

In view of relating strengths and weaknesses predicted by SWOTein to large conformational changes in protein structures, we applied SWOTein to the lysine/arginine/ornithine-binding protein from *Salmonella typhimurium*. This protein is composed of two lobes that are connected by two short hinges (hinge I: residues 89-92 and hinge II: 186-194). Two stable conformations are known: the open form in which the two lobes do not interact (X-ray structure 2LAO) and the closed form in which they do (X-ray structure 1LST) [35]. The closed form is obtained upon binding of a Lys ligand to the open form, which initiates a conformational change corresponding basically to a rigid body movement of one lobe with respect to the other. This conformational change is mainly induced by rotations of connecting hinge residues which modify their backbone torsion angle values. The two forms are shown in Fig. 7.

**Figure 7.**
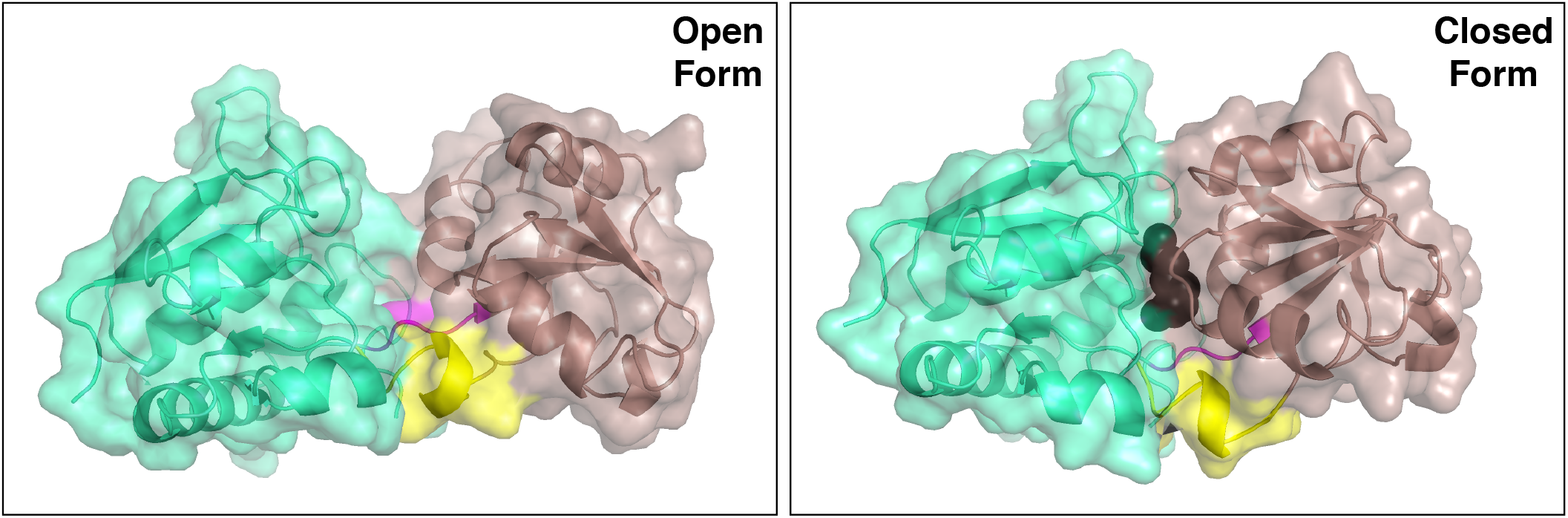
3D structure of the two stable conformations of the lysine/arginine/ornithine-binding protein. (a) Open form (PDB code 2LAO); (b) closed form (PDB code 1LST). The first lobe of the protein is colored in green, the second lobe in brown and the two hinges in magenta (hinge I) and yellow (hinge II). The Lys ligand is depicted in black spheres in the closed form.

We applied SWOTein to calculate the folding free energy contributions of each residue in the open and closed conformations. The ten weakest residues in each form, according to 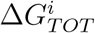, and their per-residue folding free energy contributions are listed in Table 4. Interestingly, three of the four residues of hinge I (90-92) are among the 10 weakest residues in both forms, especially when computed from the torsion angle and solvent accessibility potentials. The fourth residue (89) is among the 10 weakest residues in the closed form and the 13th weakest residue in the open form. Several residues that are close to hinge I along the sequence (84, 86, 93) or in space (160-161) are also among the 10 weakest residues in one or both forms. Even though they are not in the hinge, they are probably important for the conformational movement. This leads us to define an extended hinge I, containing residues 84-93.

**Table 4:**
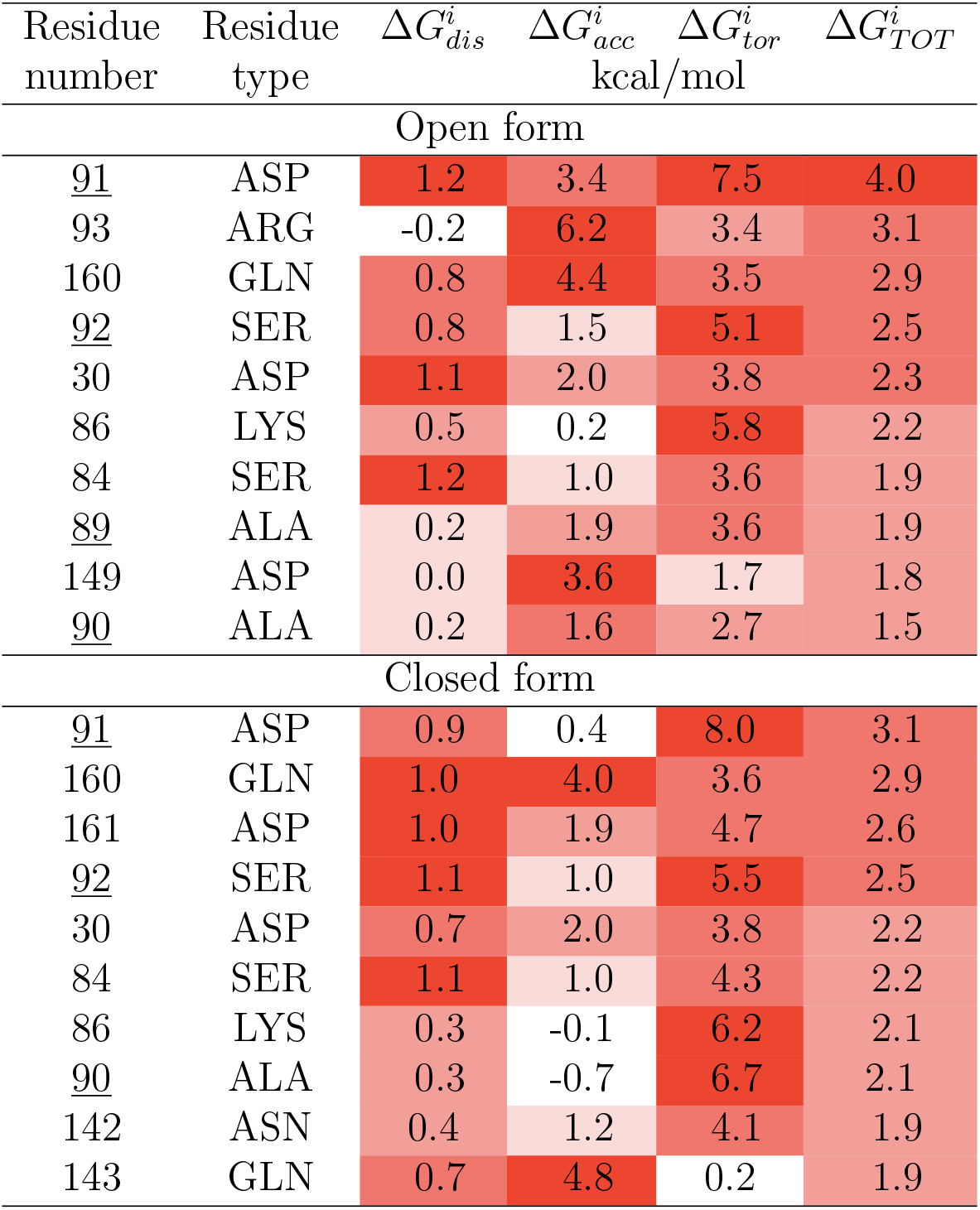
Ten weakest residues predicted by SWOTein in the open form (2LAO) and closed form (1LST) of the lysine/arginine/ornithine-binding protein. The ranking follows 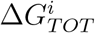. The color code is defined in Table 2. The residues that belong to hinge I are underlined.

In contrast, hinge II is quite stable, with some residues predicted as strengths and the others as neutral. This hinge undergoes movements upon the conformational change, even though to a lesser extent than hinge I [35].

The folding free energy contributions averaged over the residues in hinge II and over the residues in the extended hinge I are shown in Table 5. The extended hinge I is a weak region for the solvent accessibility and torsion angle potentials and hinge II is a strength for the torsion angle potential. It is thus clearly the extended hinge I region which drives the conformational change.

**Table 5:**
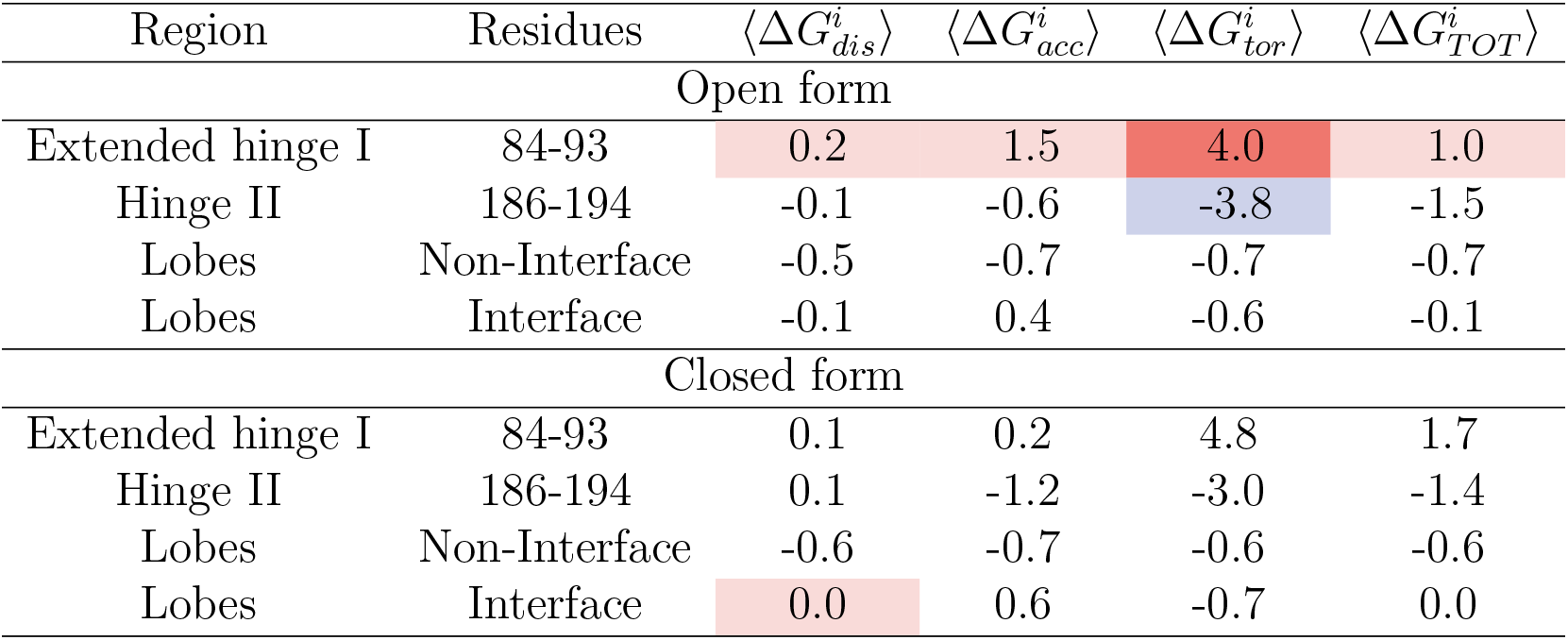
Average folding free energy contributions (in kcal/mol) for the different regions of the lysine/arginine/ornithine-binding protein. The open-closed interface between the lobes contains the residues that interact in the closed form but not in the open form, i.e. residues 1-83,94-185,195-238; they are defined as the residues whose solvent accessibility changes by more than 5% upon conformational change; the non-interface region contains the other residues. The color code is defined in Table 2.

Moreover, the residues of the two lobes that form the open-closed interface, defined as the residues that are not in contact in the open form but are in contact in the closed form, are on the average weaker than the residues in the other parts of the lobes, as shown in Table 5. This is not surprising as the they must form and break upon conformational change and must not be too strong. Stabilizing them would make the conformational change more difficult.

Finally, the residues that closely interact with the Lys ligand in the closed form, *i.e*. Tyr 14, Phe 52 and Leu 117, are weak residues in the open form for the solvent accessibility potential 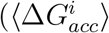 = 1.8 kcal/mol). As these residues have a functional role, which involves partial unfolding to let the ligand enter inside the structure, it is thus not surprising that they are not optimized for stability. Upon ligand binding, the solvent accessibility of these residues is reduced: they are partially buried in the open form and become completely buried in the closed form. Concomitantly, they become strengths, with 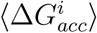 = −0.63 kcal/mol. Note that these residues are neutral or even strengths for the other two potentials.

In summary, the analysis of the strengths and weaknesses predicted by SWOTein in the open and closed forms of the lysine/arginine/ornithine-binding protein led us to detect regions that have important roles in the conformational change and in protein-ligand interactions.

#### Subtilisin Carlsberg-Eglin c interaction

To illustrate how SWOTein can provide information on molecular recognition and protein-protein interactions, we used it to study the binding between the serine protease Subtilisin Carlsberg from *Bacillus subtilis* and the protease inhibitor Eglin c from *Hirudo medicinalis* [36]. The experimental 3D structures of the unbound proteins (PBD codes 1SBC and 1EGL) and of their complex (PDB code 1CSE) are both available. We defined the protein-protein interface as the set of residues whose solvent accessibility changes by more than 5% upon binding; this is done by considering the two monomers in 1CSE either together or separately. The resulting interface region essentially involves the reactive site loop of eglin c and the active site cleft of subtilisin.

We computed the average per-residue folding free energy 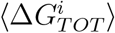 for interface residues, surface residues (accessibility>20%), and buried residues (accessibility≤20%). These energies are computed in both the unbound monomeric structures 1SBC and 1EGL (Fig. 8.a) and their complex 1CSE (Fig. 8.b). We clearly see from these histograms that the interface residues have the highest average per-site folding free energy and the buried residues the lowest. This shows, as expected, the high stability of the protein core and the lower stability of the surface, especially in interface regions. Moreover, there is a substantial gain in stability of the interface upon binding (Fig. 8.b), which is likely to be a general characteristics of protein-protein interfaces [37].

**Figure 8:**
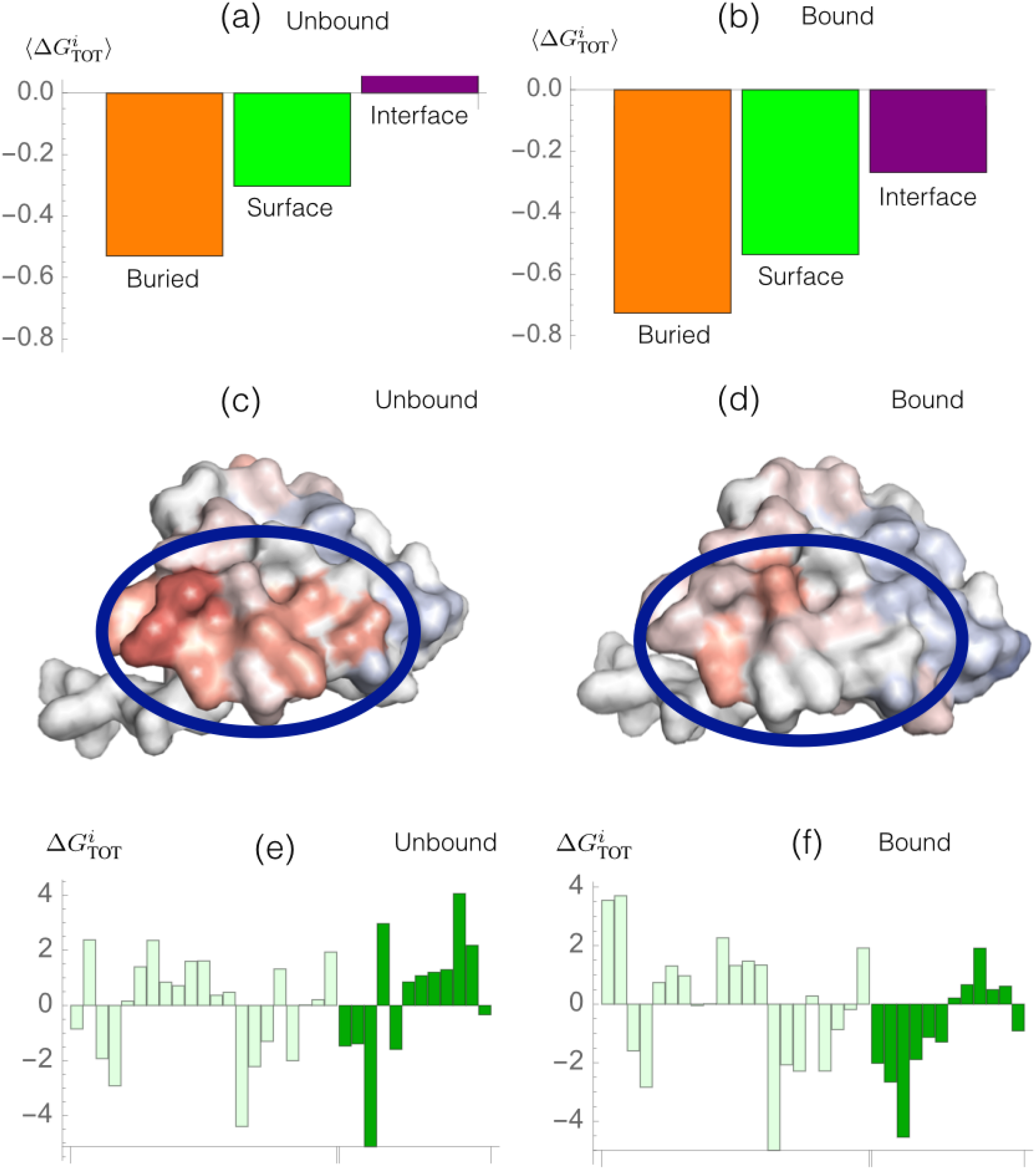
Strengths and weakness patterns in the subtilisin Carlsberg-eglin c complex predicted using the per-site folding free energies 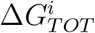. (a)-(b) Average 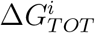 for buried, surface and interface residues in the monomeric forms (a) (PDB codes 1SBC and 1EGL) and the bound form (b) (PDB code 1CSE). (c)-(d) 3D representation of the weak and strong residues in the unbound and bound forms of eglin c (1EDL and 1CSE), using the color code defined in Table 2 (average color over the three per-site folding free energy contributions). The blue circle identifies the interface region. (e)-(f) 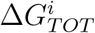 values for the interface residues in the unbound (e) and bound forms (f); light green and dark green bars represent the interface residues (in sequential order) belonging to subtilisin and eglin c, respectively.

The stabilization effect occurring at the interface is shown in detail in Fig 8.c-d where the interface residues of eglin c are colored accordingly to their stability properties in their unbound and bound forms. This region, in which most residues are weaknesses in the unbound form, gets highly stabilized upon binding with the interacting partner. Note that this strong stabilization effect is not observed in subtilisin, where almost no overall change in stability upon binding is observed (Figs 8.e-f). In this case, the stability of the complex with respect to the individual partners is essentially due to one of the two partners.

SWOTein tells us more about the way in which interacting proteins get stabilized. Indeed, a detailed per-residue analysis shows interesting patterns of stability strengths and weaknesses at the interface, as shown in Figs 8.e-f. It indicates residues whose stability is increased or decreased upon binding, and is a useful instrument for the rational modification of interacting proteins in order to stabilize or destabilize their unbound or bound conformations.

These results underline the important role of the strength/weakness patterns predicted by SWOTein in the analysis of protein-protein complex formation and of biomolecular recognition processes.

## 4 Conclusion

We introduced SWOTein and its easy-to-use webserver, devoted to predict protein residues or regions corresponding to stability strengths and weaknesses, which are often the center of crucial structural, energetic or functional characteristics. An accurate knowledge of these properties is of utmost importance in a large range of biomedical and biotechnological applications, which require understanding protein processes such as folding, molecular recognition and conformational changes. The three test cases analyzed in this paper perfectly illustrate the power of the approach. Moreover, strengths and weaknesses are interesting sites to mutate in view of modulating stability and function.

The SWOTein webserver requires as sole input the experimental or modeled protein 3D structure of the target protein. It outputs a list of per-residue folding free energy contributions and a simple visualization tool showing the strengths and weaknesses on the 3D structure (see Fig. 6.b).

One main advantage of SWOTein is that it provides, for each residue, three different contributions to the folding free energy, computed from three potentials and associated to the degree of stability of three different conformational descriptors: solvent accessibility, backbone torsion angles and inter-residue distances. The picture that comes out is complex (see *e.g*., Fig. 4): regions that are strengths for one type of potentials can be weaknesses for another one. For example, local interactions along the chain, which define secondary structures, can be strongly frustrated in a given protein region, while tertiary contacts in the same region can be quite optimal, or *vice versa*. It is clearly highly instructive to consider these contributions separately. Alternatively, if a user is only interested in overall effects, he can consider the sum of contributions, Δ*G_TOT_*.

The second advantage of SWOTein is its extreme speed. Indeed, it is able to identify strengths and weaknesses of a medium-size protein in a few tens of seconds. It can thus be applied on a large scale, even on a proteome-scale, to analyze global protein features. This will be the subject of a forthcoming study.

We will also further examine on a large scale to what extent the different kinds of per-residue folding free energy contributions correspond to different kinds of strengths and weaknesses, in order to better understand their significance and roles in the folding process, but also in other processes such as catalytic activity and allosteric mechanisms.

Other future work will aim to better understand the relation between strengths and weaknesses computed by SWOTein and strengths and weaknesses defined in the different manners described in the Introduction. We already successfully compared them with the mutational frustration index predicted by the Frustratometer [7] and are particularly curious to compare them with mutational strengths and weaknesses that incorporate the effect of natural evolution, evaluated using PoPMuSiC [12]. Not only the comparison but also the integration of the different definitions are expected to improve our insights into the structure-, dynamics- and function-related protein mechanisms.

## Acknowledgements

We thank Marie De Laet and Dimitri Gilis for their input in the early stages of the study. F.P. is postdoctoral researcher and M.R. research director at the F.R.S.-FNRS Fund for Scientific Research (Belgium); R.B. and Q.H. have previously benefitted from postdoctoral grants from the same Fund (PDR F.R.S.-FNRS project).

